# Desikan-Killiany-Tourville Atlas Compatible Version of M-CRIB Neonatal Parcellated Whole Brain Atlas: The M-CRIB 2.0

**DOI:** 10.1101/409045

**Authors:** Bonnie Alexander, Wai Yen Loh, Lillian G. Matthews, Andrea L. Murray, Chris Adamson, Richard Beare, Jian Chen, Claire E. Kelly, Peter J. Anderson, Lex W. Doyle, Alicia J. Spittle, Jeanie L.Y. Cheong, Marc L. Seal, Deanne K. Thompson

## Abstract

Our recently published M-CRIB atlas comprises 100 neonatal brain regions including 68 compatible with the widely-used Desikan-Killiany adult cortical atlas. A successor to the Desikan-Killiany atlas is the Desikan-Killiany-Tourville atlas, in which some regions with unclear boundaries were removed, and many existing boundaries were revised to conform to clearer landmarks in sulcal fundi. Our first aim here was to modify cortical M-CRIB regions to comply with the Desikan-Killiany-Tourville protocol, in order to offer: a) compatibility with this adult cortical atlas, b) greater labelling accuracy due to clearer landmarks, and c) optimisation of cortical regions for integration with surface-based infant parcellation pipelines. Secondly, we aimed to update subcortical regions in order to offer greater compatibility with subcortical segmentations produced in FreeSurfer. Data utilized were the T2-weighted MRI scans in our M-CRIB atlas, for ten healthy neonates (postmenstrual age at MRI 40-43 weeks, 4 female), and corresponding parcellated images. Edits were performed on the parcellated images in volume space using ITK-SNAP. Cortical updates included deletion of frontal and temporal poles and ‘Banks STS’, and modification of boundaries of many other regions. Changes to subcortical regions included the addition of ‘ventral diencephalon’, and deletion of ‘subcortical matter’ labels. A detailed updated parcellation protocol was produced. The resulting whole-brain M-CRIB 2.0 atlas comprises 94 regions altogether. This atlas provides comparability with adult Desikan-Killiany-Tourville-labelled cortical data and FreeSurfer-labelled subcortical data, and is more readily adaptable for incorporation into surface-based neonatal parcellation pipelines. As such, it offers the ability to help facilitate a broad range of investigations into brain structure and function both at the neonatal time point and developmentally across the lifespan.

## 1 Introduction

We recently published the M-CRIB (Alexander et al., 2017) neonatal parcellated brain atlas, comprising 100 regions in total, including 68 compatible with the Desikan-Killiany (DK; Desikan et al., 2006) adult cortical atlas, as well as basal ganglia, thalamus, and cerebellar regions. The DK atlas is one of the most commonly used parcellation schemes, thus an advantage of the M-CRIB atlas is that it provides compatibility of parcellated cortical regions between neonatal and later time points. This can help facilitate investigations into regional brain structure and function across the lifespan, potentially longitudinally. The M-CRIB atlas is also valuable in that it comprises ten individual high-quality detailed manual parcellations based on high resolution T2-weighted images, providing a combination of detailed whole-brain ‘ground truth’ and individual variability in morphology not available previously. We have recently demonstrated the applicability of the M-CRIB atlas, reporting differences in neonatal regional brain volumes based on premature birth (Alexander et al., 2018).

A successor to the DK atlas is the Desikan-Killiany-Tourville (DKT; Klein & Tourville, 2012) adult cortical parcellated atlas, in which some regions with unclear or arbitrary boundaries were removed, and many existing boundaries were revised to conform to sulcal fundi. This provides greater anatomical consistency across individuals due to clearer and more reproducible landmarks. The use of sulcal-based landmarks also optimises utility for application using surface-based labelling such as is performed in FreeSurfer (Fischl et al., 2002). Surface-based methods incorporate surface-based registration which aligns sulci and gyri more precisely than volume-based methods (Fischl, Sereno, Tootell, & Dale, 1999; Makropoulos et al., 2018), thus facilitating more precise alignment of sulcally-bounded labels.

Multiple surface-based tools have been developed for infant data (e.g., Hill et al., 2010; Kim et al., 2016; Li, Wang, Gilmore, Lin, & Shen, 2015; Li, Wang, Shi, Lin, & Shen, 2014; Makropoulos et al., 2018). FreeSurfer tools for infant parcellation are currently in development (e.g., Zollei, Ou, Iglesias, Grant, & Fischl, 2017). A key resource facilitating accurate surface-based parcellation at the neonatal time point is high-quality ground truth neonatal parcellated training data. Such data are currently in strong demand.

Here we firstly aimed to modify the cortical regions and protocol of the existing volumetric M-CRIB atlas to comply with the DKT cortical parcellation protocol, in order to a) offer compatibility with data at older time points parcellated with the adult DKT atlas, b) achieve greater anatomical consistency in labelling across brains due to some boundaries being revised to clearer landmarks in sulcal fundi, and c) offer greater ease of adaptability for integration into neonatal surface-based parcellation pipelines due to the use of these sulcally-defined boundaries. Secondly, we aimed to update subcortical regions to offer greater compatibility with those segmented by FreeSurfer’s subcortical pipeline, including addition of the ‘ventral diencephalon’, and removal of ‘subcortical matter’ labels. These cortical and subcortical updates together comprise the ‘M-CRIB 2.0’ neonatal atlas.

## 2 Methods

### 2.1 Data

Data utilized were the individual segmentation images and T2- and T1-weighted images comprising the M-CRIB atlas. This sample consisted of 10 healthy term-born (≥37 weeks’ gestation) neonates (4 female, 6 male; gestational age at scanning 40.29– 43.00 weeks, M = 41.71, SD = 1.31), selected from a larger cohort of controls with MRI scans recruited as part of preterm studies (Spittle et al., 2014; Walsh, Doyle, Anderson, Lee, & Cheong, 2014). T2-weighted images were acquired using a transverse T2 restore turbo spin echo sequence with: 1 mm axial slices, flip angle = 120°, TR = 8910 ms, TE = 152 ms, FOV = 192 × 192 mm, in-plane resolution 1 mm^2^ (zero-filled interpolated to 0.5 × 0.5 × 1 mm), matrix = 384 × 384. Three-dimensional T1-weighted images were acquired using a magnetisation prepared rapid gradient-echo sequence with: 1 mm axial slices, flip angle = 9°, TR = 2100 ms, TE = 3.39 ms, FOV = 192 × 192 mm, in-plane resolution 1 mm^2^ (zero-filled interpolated to 0.5 × 0.5 × 1 mm), matrix = 384 × 384. T2-weighted images were bias-corrected using N4ITK (Tustison et al., 2010), skull-stripped using BET (Smith, 2002; Smith et al., 2004), aligned to the anterior commissure-posterior commissure axis with 3D Slicer v.4.1.1 (http://www.slicer.org/) (Fedorov et al., 2012), and resampled to 0.63 × 0.63 × 0.63 mm isotropic voxels (preserving voxel volume) using FLIRT (Greve & Fischl, 2009; Jenkinson, Bannister, Brady, & Smith, 2002; Jenkinson & Smith, 2001). Further information about the sample, data and preprocessing is listed in Alexander et al. (2017).

T1-weighted images are included in the M-CRIB and M-CRIB 2.0 datasets, however they were not used for manual tracing, because of low contrast between tissue types due to partial myelination at the neonatal time point. Rather, they are included as they may provide additional intensity information leverageable in multimodal automated parcellation pipelines. The T2-weighted images, which confer higher tissue contrast, were used both for parcellation of the original M-CRIB, and for the edits performed here.

### 2.2 Manual editing procedure

The individual segmentation images comprising the M-CRIB atlas were edited in volume space using Insight Toolkit (ITK)-SNAP v3.6.0 (itksnap.org) (Yushkevich et al., 2006), by one operator (B.A.). ITK-SNAP displays axial, sagittal, and coronal views and a composite 3D mesh representation of utilised labels. The edits were performed and checked on a combination of the axial, sagittal, and coronal views, with reference to the 3D surface view. Edits were performed region-by-region rather than brain-by-brain, except in some areas where edits to multiple adjacent regions were required, as the alterations to one region sometimes necessitated specification of adjacent areas’ boundaries. For some regions such as the newly-specified ventral diencephalon, edits were performed for the whole sample, and then checked and edited where necessary to ensure consistency.

### 2.3 Parcellation protocols

In the following cortical protocol, revised boundary descriptors are listed that aimed to replicate the DKT (Klein & Tourville, 2012) protocol as closely as possible within this volumetric neonatal sample. Where possible, verbatim DKT boundary descriptors have been utilised, and are indicated in bold font. Descriptors retained from the DK protocol are indicated in italics. Descriptors either retained from the M-CRIB protocol or newly specified here are indicated in regular font. Some anatomical axis descriptors (e.g., ‘anterior’) have been adjusted to retain anatomical accuracy in volume space.

When revising the boundary descriptors and editing the data, reference was made to Klein and Tourville (2012), the anatomical atlas by Petrides (2011) which describes many sulci used as DKT boundaries, other anatomical atlases (Duvernoy, Bourgouin, Cabanis, & Cattin, 1999; Duvernoy, 2013; Rubin & Safdieh, 2007), the BrainInfo database (National Primate Research Center, 1991-present; www.braininfo.org), and individual papers describing anatomy (Dumoulin et al., 2000; Nagata, Rhoton, & Barry, 1988; Türe, Yasargil, Al-Mefty, & Yasargil, 1999; Watson et al., 1993).

Updates between the M-CRIB and M-CRIB 2.0 atlases pertain to the DKT cortical regions, ventral diencephalon (added), brainstem (edited in the course of defining ventral diencephalon), left and right ‘subcortical matter’ (removed), and left and right cerebral white matter (edited in removal of subcortical matter labels). Cerebellum, hippocampus, amygdala, and ventricles, were retained as per the original M-CRIB atlas, and parcellation protocols for these regions are listed in Alexander et al. (2017). Basal ganglia and thalamus were not manually edited and protocols for these regions are retained from the M-CRIB atlas, however postprocessing performed on these segmentations was removed, as described below.

### 2.4 M-CRIB 2.0 protocol

#### 2.4.1 Cortical regions

Frontal pole, temporal pole, and “banks of the superior temporal sulcus” regions were removed as per the DKT protocol, and replaced with surrounding gyral labels.

##### 2.4.1.1 Temporal - medial aspect

***Entorhinal cortex*** *<u>Boundaries: Anterior:</u>* **Temporal incisure (rostral limit of collateral sulcus).** *<u>Posterior:</u>* **Posterior limit of the amygdala.** Superior: **Medio-dorsal margin of the temporal lobe anteriorly, amygdala** and hippocampus **posteriorly.** *<u>Medial:</u> Medial aspect of the temporal lobe. <u>Lateral:</u>* **Rhinal sulcus (collateral sulcus), or the collateral sulcus if the rhinal sulcus is not present.**

***Parahippocampal gyrus*** *<u>Boundaries: Anterior:</u>* **Posterior limit of the amygdala.** *<u>Posterior:</u>* **Posterior limit of the hippocampus.** *<u>Medial:</u> Medial aspect of the temporal lobe. <u>Lateral:</u> Collateral sulcus.*

***Temporal pole (removed)* The area included in the DK temporal pole has been redistributed to the superior, middle and inferior temporal gyrus regions.**

***Fusiform Gyrus*** *<u>Boundaries: Anterior:</u>* **Anterior limit of occipitotemporal sulcus (anterior limit of collateral sulcus).** *<u>Posterior:</u>* **First transverse sulcus posterior to the temporo-occipital notch.** This is consistent with the posterior extent of the existing parcellation, which was based on the M-CRIB boundary listed as “posterior transverse collateral sulcus (Duvernoy et al., 1999).” *<u>Medial:</u>* **Collateral sulcus.** *<u>Lateral:</u>* **Occipitotemporal sulcus.**

##### 2.4.1.2 Temporal – lateral aspect

***Superior temporal gyrus*** *<u>Boundaries: Anterior:</u>* **Anterior limit of the superior temporal sulcus or a projection from the superior temporal sulcus to the anterior limit of the temporal lobe.** *<u>Posterior:</u>* **Junction of posterior horizontal ramus of the lateral sulcus (or its posterior projection) and caudal superior temporal sulcus (**1^st^**segment of the caudal superior temporal sulcus).** Note: The DKT protocol lists 1^st^, 2^nd^ or 3^rd^ segment, however the current parcellations of this region posteriorly conform specifically to the landmark that Petrides (2011) describes as the 1^st^ segment, i.e., bounding the posterior extent of supramarginal gyrus (Petrides, 2011). *<u>Superomedial:</u> Lateral fissure (and when present, the supramarginal gyrus* and insula) *<u>Inferior:</u>* **Superior temporal sulcus.**

***Middle temporal gyrus*** *<u>Boundaries: Anterior:</u>* **Anterior limit of the superior temporal sulcus.** *<u>Posterior:</u>* **Anterior occipital sulcus.** Note: this has also been described as the ascending limb of inferior temporal sulcus (Dumoulin et al., 2000; Petrides, 2011; Watson et al., 1993). This is described by Duvernoy et al. (1999) as only sometimes being present: “The inferior temporal sulcus is usually not continuous and does not provide easy identification. In the vicinity of the occipital lobe, its posterior end may occasionally run upward and be called the anterior occipital sulcus.” In cases where this sulcus segment did not occur, the boundary was a point on a theoretical line extending vertically from the *occipito-temporal incisure on the cortical surface. <u>Superomedial:</u>* **Superior temporal sulcus anteriorly, posteriorly formed by caudal superior temporal sulcus third segment.** *<u>Inferior:</u>* **Inferior temporal sulcus.**

***Inferior temporal gyrus*** *<u>Boundaries: Anterior:</u>* **Anterior limit of the inferior temporal sulcus.** *<u>Posterior:</u>* **Anterior occipital sulcus** (see descriptor for posterior boundary of middle temporal gyrus). In cases where this sulcus segment did not occur, the boundary was a point on a theoretical line extending vertically from the occipito-temporal incisure on the cortical surface. *<u>Superior:</u>* Inferior temporal sulcus *<u>Inferior:</u>* Occipitotemporal sulcus (Duvernoy et al., 1999).

***Transverse temporal cortex*** *<u>Description:</u>* Also termed Heschl’s gyrus, this area lies along the superior temporal plane, extending from the retroinsular region to the lateral edge of the superior temporal gyrus. It can be a single gyrus, or divided into two gyri by an intermediate transverse temporal sulcus (Duvernoy et al., 1999; Rademacher, 2003). *<u>Boundaries: Anterior:</u>* **Anterior limit of first transverse temporal sulcus** (also referred to as the anterior transverse temporal sulcus (Tamraz & Comair, 2006).) *<u>Posterior:</u>* **Posterior limit of Heschl’s sulcus** (also referred to as the posterior transverse temporal sulcus (Rademacher, 2003; Tamraz & Comair, 2006), or transverse temporal sulcus (Duvernoy et al., 1999; Ono, Kubik, & Abernathey, 1990)). *<u>Medial:</u>* Retro-insular area of the lateral fossa. *<u>Lateral:</u>* Lateral surface of the superior temporal gyrus.

##### 2.4.1.3 Frontal

***Superior frontal gyrus*** *<u>Boundaries: Anterior:</u>* **Frontomarginal sulcus.** *<u>Posterior:</u>* **Precentral sulcus (lateral surface); paracentral sulcus (medial surface).** *<u>Medial:</u> Medial aspect of the frontal lobe. <u>Inferior:</u> Superior frontal sulcus.*

***Middle frontal gyrus - rostral division*** *<u>Description:</u>* Approximates the rostral-most three quarters of the middle frontal gyrus. *<u>Boundaries: Anterior:</u>* **Anterior limit of the superior frontal sulcus.** *<u>Posterior:</u>* A theoretical line separating the caudal-most quarter of the middle frontal gyrus. *<u>Medial:</u>* **Superior frontal sulcus.** *<u>Lateral:</u>* **Inferior frontal sulcus; anterior to inferior frontal sulcus, the ventro-lateral boundary is formed by frontomarginal sulcus and lateral H-shaped orbital sulcus.**

***Middle frontal gyrus - caudal division*** *<u>Description:</u>* Approximates the caudal-most quarter of the middle frontal gyrus. *<u>Boundaries: Anterior:</u>* A theoretical line separating the caudal-most quarter of the middle frontal gyrus. *<u>Posterior:</u>* **Precentral sulcus.** *<u>Medial:</u>* **Superior frontal sulcus.** *<u>Lateral:</u>* **Inferior frontal sulcus.**

***Inferior frontal gyrus*** *<u>Description:</u>* The inferior frontal gyrus comprises the three pars regions.

***Inferior frontal gyrus - pars opercularis*** *<u>Boundaries: Anterior:</u>* **Anterior ascending ramus of the lateral sulcus,** which is also referred to as the ascending ramus (Türe et al., 1999) (Türe, Yasargil, Al-Mefty, & Yasargil, 1999). *<u>Posterior:</u>* **Precentral sulcus.** *<u>Superomedial:</u>* **Inferior frontal sulcus.** *<u>Inferomedial:</u>* **Circular insular sulcus.**

***Inferior frontal gyrus - pars triangularis*** *<u>Boundaries: Anterior:</u>* **Pretriangular sulcus.** *<u>Posterior:</u>* **Anterior ascending ramus of the lateral sulcus.** *<u>Superomedial:</u>* **Inferior frontal sulcus.** *<u>Inferomedial:</u>* **Anterior horizontal ramus of the lateral sulcus; if the anterior horizontal ramus of the lateral sulcus does not extend anteriorly to pretriangular sulcus, an anterior projection from anterior horizontal ramus of the lateral sulcus to pretriangular sulcus.**

***Inferior frontal gyrus - pars orbitalis*** *<u>Boundaries: Anterior:</u>* **Pretriangular sulcus – if pretriangular sulcus does not extend ventrally to the lateral H-shaped orbital sulcus, a ventral projection from pretriangular sulcus to lateral H-shaped orbital sulcus completes the anterior boundary.** *<u>Posterior:</u>* **Posterior limit of orbitofrontal cortex.** *<u>Superomedial:</u>* **Anterior horizontal ramus of the lateral sulcus – if the anterior horizontal ramus of the lateral sulcus does not extend anteriorly to the pretriangular sulcus, an anterior projection from anterior horizontal ramus of the lateral sulcus to pretriangular sulcus completes the lateral boundary.** *<u>Inferomedial:</u>* **Lateral H-shaped orbital sulcus.**

***Orbitofrontal cortex - lateral division*** *<u>Boundaries: Anterior:</u>* **Frontomarginal sulcus.** *<u>Posterior:</u>* **Posterior limit of orbitofrontal cortex.** *<u>Medial:</u>* **Olfactory sulcus.** *<u>Lateral:</u>* **Lateral H-shaped orbital sulcus.**

***Orbitofrontal cortex - medial division*** *<u>Boundaries: Anterior:</u>* **Frontomarginal sulcus.** *<u>Posterior:</u>* **Posterior limit of orbitofrontal cortex.** *<u>Superomedial:</u>* **superior rostral sulcus; if superior rostral sulcus merges with cingulate sulcus, the medial/dorsal boundary is formed by cingulate sulcus.** *<u>InferoLateral:</u>* **Olfactory sulcus.**

***Precentral gyrus*** *<u>Boundaries: Anterior:</u>* **Precentral sulcus.** *<u>Posterior:</u>* **Central sulcus.** *<u>Superomedial:</u>* Medial bank of the central sulcus. *<u>Inferomedial:</u>* **Circular insular sulcus.**

***Paracentral lobule*** *<u>Description:</u>* Medial structure consisting of the superomedial ends of the precentral and postcentral gyri surrounding the superior end of the central sulcus (Duvernoy et al., 1999). *<u>Boundaries: Anterior:</u>* **Paracentral sulcus.** *<u>Posterior:</u>* **Marginal ramus of cingulate sulcus.** *<u>Inferomedial:</u>* **Cingulate sulcus.** *<u>Superolateral:</u>* Medial bank of the central sulcus.

##### 2.4.1.4 Parietal

***Postcentral gyrus*** *<u>Boundaries: Anterior:</u>* **Central sulcus.** *<u>Posterior:</u>* **Postcentral sulcus.** *<u>Superomedial:</u>* Medial bank of the central sulcus. *<u>Inferomedial:</u>* **Circular insular sulcus – if the lateral limit of postcentral sulcus extends anterior to circular insular sulcus, the posterior portion of the lateral/ventral boundary is formed by the lateral sulcus.**

***Supramarginal gyrus*** *<u>Description:</u>* **Formed by sulci demarcating the cortical convolution surrounding the posterior ascending ramus of the lateral sulcus.** *<u>Boundaries: Anterior:</u>* **Postcentral sulcus.** *<u>Posterior:</u>* **Primary intermediate sulcus** supero**medially, and caudal superior temporal sulcus (first segment)** infero**laterally.** *<u>Superomedial:</u>* **Intraparietal sulcus.** *<u>Inferior:</u>* **Lateral sulcus anterior to posterior horizontal ramus of the lateral sulcus, posterior horizontal ramus of the lateral sulcus posteriorly.**

***Superior parietal cortex*** *<u>Boundaries: Anterior:</u>* **Postcentral sulcus.** *<u>Posterior:</u>* **Transverse sulcus lying immediately posterior to the parietooccipital sulcus – this is described as the transverse occipital sulcus, medial segment, by Petrides (2011).** *<u>Medial:</u>* **Dorsomedial hemispheric margin.** *<u>Lateral:</u>* **Intraparietal sulcus.**

***Inferior parietal cortex*** *<u>Description:</u> Includes the inferior parietal gyrus and the angular gyrus and lies inferior to the superior parietal gyrus. <u>Boundaries: Anterior:</u>* **Caudal superior temporal sulcus, first segment.** *<u>Posterior:</u>* A theoretical line reaching from the parieto-occipital fissure to the temporo-occipital incisure. *<u>Medial:</u>* **Intraparietal sulcus.** *<u>Lateral:</u>* **lateral occipital sulcus anteriorly, transverse occipital sulcus lateral segment posteriorly.**

***Precuneus cortex*** *<u>Boundaries: Anterior:</u>* Marginal segment of the cingulate sulcus (Duvernoy et al., 1999). *<u>Posterior:</u>* **Parieto-occipital sulcus.** *<u>Inferior:</u>* Subparietal sulcus. *<u>Medial:</u>* Medial surface of the hemisphere. *<u>Lateral</u>: Superior parietal gyrus*.

##### 2.4.1.5 Occipital

***Lingual gyrus*** *<u>Boundaries: Anterior:</u>* **Posterior limit of the hippocampus.** *<u>Posterior:</u>* **Posterior limit of calcarine sulcus.** *<u>Medial:</u> Medial portion of the temporal and occipital cortices. <u>Lateral:</u>* **Collateral sulcus.**

***Pericalcarine cortex*** *<u>Boundaries: Anterior:</u>* **Junction of calcarine sulcus and parietooccipital sulcus.** *<u>Posterior:</u>* **Posterior limit of calcarine sulcus.** *<u>Superior:</u>* **Dorsomedial margin of calcarine sulcus.** *<u>Inferior:</u>* **Ventromedial margin of calcarine sulcus.** *<u>Medial:</u> Medial portion of the temporal and occipital cortices. <u>Lateral:</u>* The depth of the calcarine sulcus.

***Cuneus cortex*** *<u>Boundaries: Anterior:</u>* **Parietooccipital sulcus.** *<u>Posterior:</u>* **Posterior limit of calcarine sulcus.** *<u>Ventral:</u>* **Dorsomedial margin of calcarine sulcus.** *<u>Dorsal:</u>* **Dorsomedial hemispheric margin.**

***Lateral occipital cortex*** *<u>Boundaries: Anterior:</u>* **Temporo-occipital notch laterally, anterior occipital sulcus more medially, transverse occipital sulcus, medial segment, medial to intraparietal sulcus.** *<u>Posterior:</u> The last visible portion of occipital cortex. <u>Medial:</u> Cuneus/pericalcarine cortex. <u>Lateral:</u>* The lateral surface of the hemisphere at this area’s anterolateral boundaries.

##### 2.4.1.6 Cingulate

***Rostral anterior division*** *<u>Boundaries: Anterior:</u>* ***Cingulate sulcus***. *<u>Posterior:</u>* ***Corpus callosum genu***. Specifically, on the sagittal plane, a theoretical line intersecting at approximately 45 degrees with the genu. See Alexander et al. (2017) for further illustration. *<u>Ventral:</u>* **Dorsal to the corpus callosum, the ventral boundary is formed by the callosal sulcus. In the subgenual area, it is formed by the cingulate sulcus.** In the case of “double parallel cingulate” sulcus that continues anteroventrally to join the ‘superior rostral sulcus’ (listed in Klein & Tourville, 2012), the ventral boundary is the superior rostral sulcus, also termed ‘supraorbital sulcus’ (Duvernoy et al., 1999, p.33).

***Caudal anterior division*** *<u>Boundaries: Anterior:</u>* **Corpus callosum genu**. *<u>Posterior:</u>* **Mammillary bodies.** *<u>Rostral/dorsal:</u>* **Cingulate sulcus; in the event of a “double parallel cingulate,” (e.g., Ono et al., 1990), the rostral/dorsal boundary of the cingulate is formed by the more rostral-dorsal branch of the cingulate sulcus.** *<u>Ventral:</u>* **Callosal sulcus.**

***Posterior division*** *<u>Boundaries: Anterior:</u>* **Mammillary bodies.** *<u>Posterior:</u>* **Junction of the subparietal sulcus and cingulate sulcus (approximately).** *<u>Superior:</u>* Cingulate sulcus. *<u>Ventral:</u>* **Callosal sulcus.**

***Isthmus division*** *<u>Boundaries: Anterior:</u>* **Junction of the subparietal sulcus and cingulate sulcus (approximately).** *<u>Posterior:</u>* The anterior calcarine sulcus (Duvernoy et al., 1999) if present, or the parieto-occipital fissure. *<u>Lateral:</u>* The depth of the calcarine sulcus.

##### 2.4.1.7 Insula

***Insula*** *<u>Description:</u>* Inverted-triangle-shaped area of mesocortex in the base of the lateral fossa covered by frontal, temporal, central and parietal opercula; and delineated from these by the circular insular sulcus (also termed periinsular or limiting sulcus) (Duvernoy et al., 1999; Türe et al., 1999). *<u>Boundaries: Anterior:</u>* Anterior peri-insular sulcus. *<u>Superior:</u>* Superior peri-insular sulcus. *<u>Infero-Posterior:</u>* Inferior peri-insular sulcus.

#### 2.4.2 Subcortical regions

##### 2.4.1.2 Basal ganglia and thalamus

The manual tracing protocol for the M-CRIB basal ganglia nuclei (caudate, putamen, pallidum and nucleus accumbens) and thalamus is described in Loh et al. (2016). These regions were not manually edited here. However, in the original M-CRIB dataset, basal ganglia and thalamus segmentations underwent morphological smoothing. For the M-CRIB 2.0, the smoothed segmentations of these structures were replaced with non-smoothed segmentations in order to recover fine-scale, irregular, intensity-based anatomical detail such as is provided for the rest of the M-CRIB and M-CRIB 2.0 regions.

##### 2.4.1.3 Ventral diencephalon

The protocol for this region is based on that of de Macedo Rodrigues et al. (2015). *<u>Boundaries: Anterior:</u>* Anterior commissure (however, unlike the protocol of de Macedo Rodrigues et al., where the infero-rostral boundary is designated as the infundibular recess, we have referred solely to the anterior commissure as an anterior boundary, as much of the optic recess was also visible posterior to the anterior commissure). *<u>Posterior:</u>* Medially, the posterior commissure. Laterally, the posterior extent of the lateral geniculate nucleus. However, the lateral geniculate nucleus itself was retained as part of the thalamus label. *<u>Superior:</u>* The inferior surface of the thalamus, posteriorly (as per de Macedo Rodrigues et al., 2015). *<u>Inferior:</u>* A line extending from the pontomesencephalic sulcus anteriorly, to the posterior commissure posteriorly. *<u>Lateral:</u>* The optic pathways (de Macedo Rodrigues et al., 2015).

##### 2.4.1.4 Brainstem

The M-CRIB brainstem label was originally derived via the initial automated MANTiS (Beare et al., 2016) tissue segmentation, and refined during the process of manually delineating surrounding structures. Here partial sections of the cerebral peduncles, red nucleus, and substantia nigra have been reassigned from the brainstem label to form part of the ventral diencephalon label.

## 3 Results

The M-CRIB 2.0 atlas comprises 94 regions: 62 cortical regions, and subcortical and cerebellar regions from the M-CRIB atlas. Figures 1 and 2 illustrate some of the updates made, displayed on surface meshes and axial slices, respectively. Atlas colours and corresponding label names are shown in Supplementary Figure S1.

**Figure 1.**
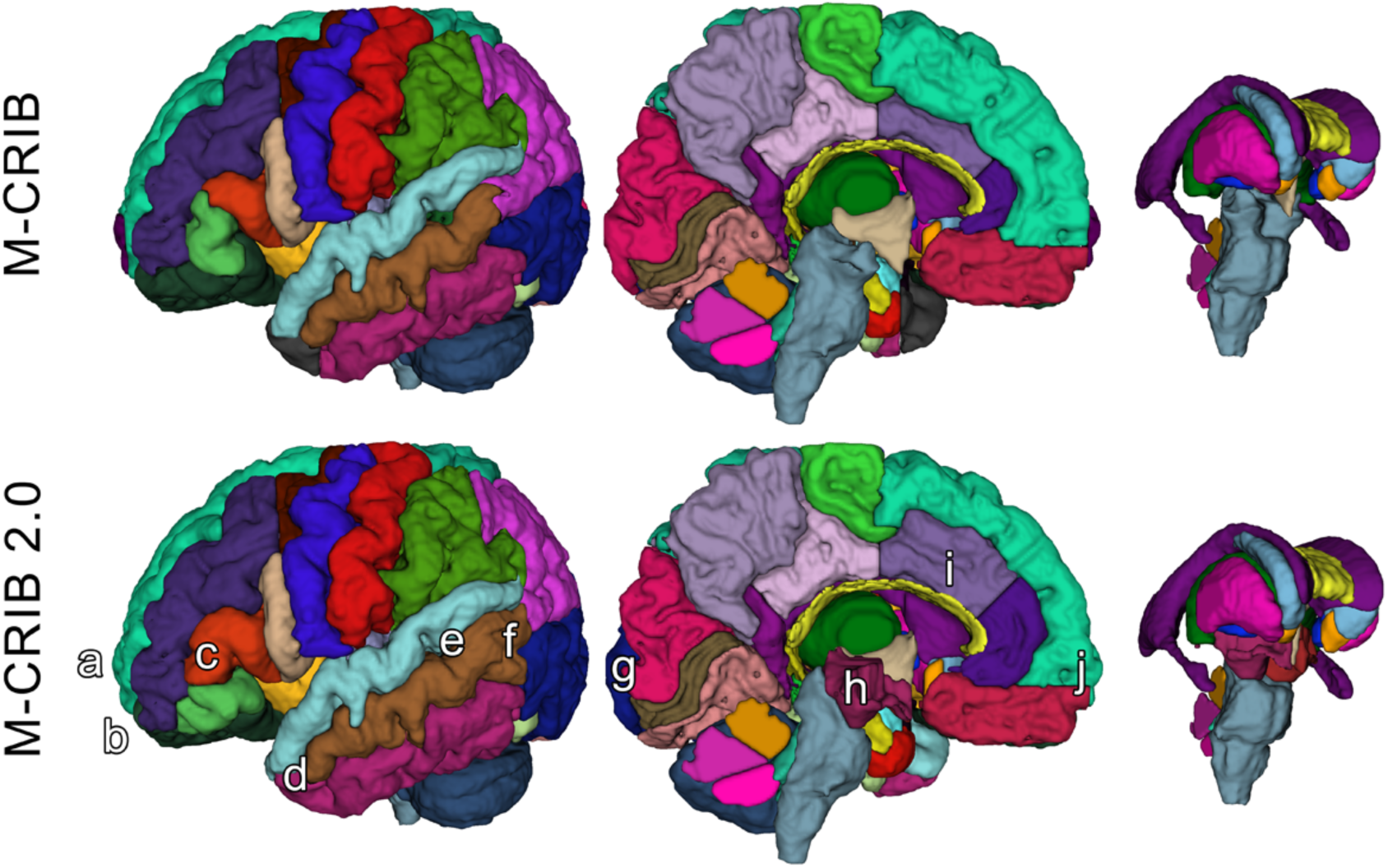
Surface meshes of a single left hemisphere (lateral and medial views) and some subcortical structures, of a single participant, illustrating some examples of updated regions. Top row: original M-CRIB atlas. Bottom row: M-CRIB 2.0. Annotations indicate some of the updates made: a) removal of frontal pole, b) revision of boundary between lateral orbitofrontal (dark green) and rostral middle frontal (dark blue) regions, c) revision of boundaries of ‘pars’ regions of inferior frontal gyrus. d) replacement of temporal pole (dark grey) with superior, middle and inferior temporal labels, e) replacement of ‘banks STS’ (dark green) region with superior and middle temporal labels, f) revision of boundary between lateral occipital (dark purple) and temporal regions, g) revision of medial boundary of lateral occipital region, h) addition of ‘ventral diencephalon’ (maroon) which replaces sections of brainstem (grey) and removed ‘subcortical matter’ (not shown) label, i) revision of rostral and caudal anterior cingulate (purple) regions to encompass cortex extending to the more rostral/dorsal branch of a parallel double cingulate sulcus.

**Figure 2.**
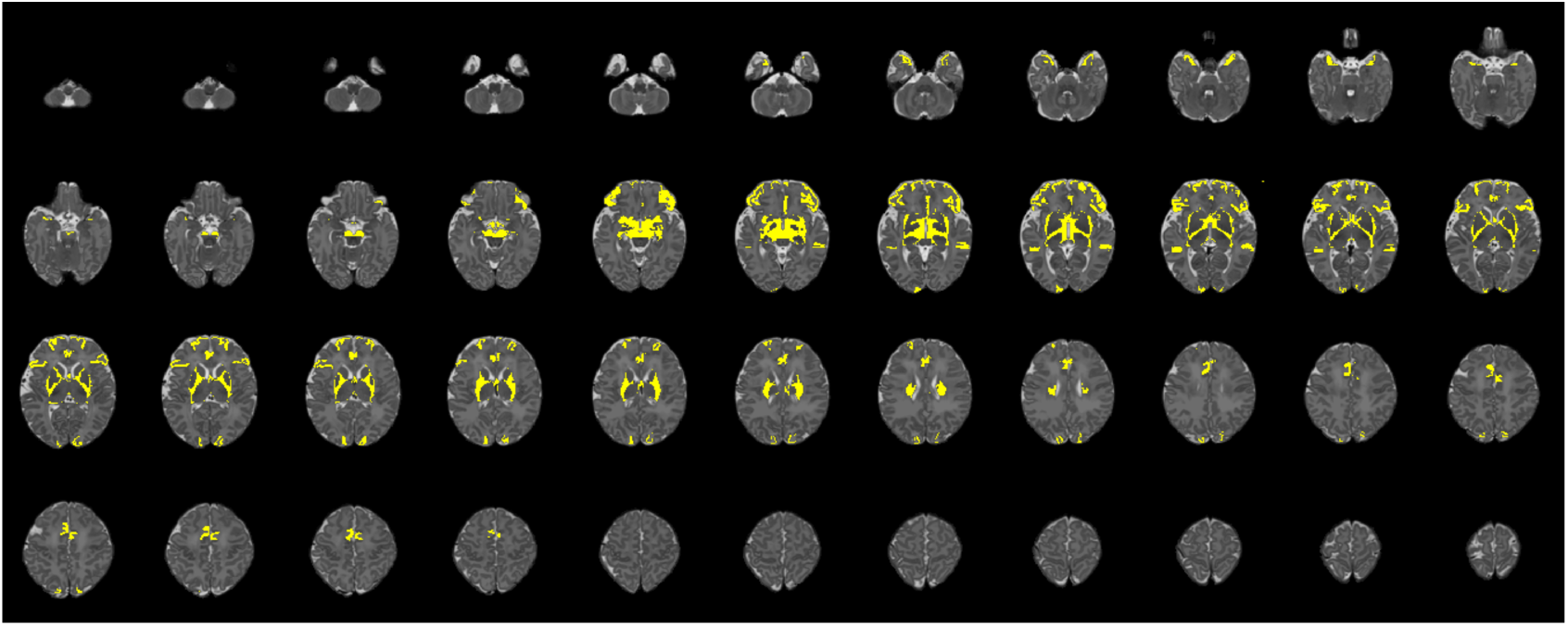
Axial slices for a single participant, illustrating regions (shown in yellow) where edits were made to update the original M-CRIB parcellated image to the M-CRIB 2.0 parcellation. Slices are presented in order from inferior (top left) to superior (bottom right), with every third slice displayed.

In Figure 2, altered regions surrounding basal ganglia and thalamus primarily reflect the removal of the ‘subcortical matter’ label, which was replaced with ‘ventral diencephalon’ and cerebral white matter labels.

Table 1 lists the mean volume of each M-CRIB 2.0 region, and the volume relative to the equivalent structure, where applicable, from the original M-CRIB atlas.

**Table 1.** Mean volume of each M-CRIB 2.0 region, and volume relative to equivalent M-CRIB region

| Region | Structure | Mean volume in mm<sup>3</sup>, (standard deviation), % volume of original M-CRIB area |  |  |  |  |  |
| --- | --- | --- | --- | --- | --- | --- | --- |
|  | CSF (extraventricular) | S 89668 | (18597) | 100 |  |  |  |
|  | brainstem | S 5394 | (632) | 85 |  |  |  |
|  | corpus callosum | S 764 | (179) | 100 |  |  |  |
|  | L/R ventral diencephalon | L 1195 | (107) | N/A | R 1216 | (121) | N/A |
|  | L/R cerebral white matter | L 79958 | (7244) | 106 | R 80610 | (6890) | 106 |
|  | L/R hippocampus | L 1473 | (157) | 100 | R 1430 | (170) | 100 |
|  | L/R amygdala | L 840 | (86) | 100 | R 802 | (91) | 100 |
|  | L/R lateral ventricle | L 2965 | (841) | 100 | R 2212 | (771) | 100 |
|  | 3rd ventricle | S 481 | (47) | 100 |  |  |  |
|  | 4th ventricle | S 410 | (70) | 100 |  |  |  |
| Cerebellum | cerebellar vermis anterior | S 214 | (34) | 100 |  |  |  |
|  | cerebellar vermis superior posterior | S 201 | (39) | 100 |  |  |  |
|  | cerebellar vermis inferior posterior | S 214 | (38) | 100 |  |  |  |
|  | L/R cerebellar hemisphere | L 11374 | (1473) | 100 | R 11390 | (1466) | 100 |
| Basal ganglia and thalamus | L/R caudate | L 1370 | (174) | 104 | R 1333 | (136) | 103 |
|  | L/R putamen | L 1757 | (142) | 103 | R 1772 | (126) | 102 |
|  | L/R accumbens area | L 189 | (38) | 115 | R 188 | (31) | 108 |
|  | L/R pallidum | L 775 | (82) | 102 | R 770 | (79) | 102 |
|  | L/R thalamus | L 3904 | (409) | 100 | R 3896 | (419) | 100 |
| Cortex | L/R unknown | L 202 | (42) | 103 | R 224 | (50) | 104 |
|  | L/R caudal anterior cingulate | L 1080 | (474) | 120 | R 1306 | (259) | 126 |
|  | L/R caudal middle frontal | L 1487 | (272) | 100 | R 1545 | (226) | 100 |
|  | L/R cuneus | L 1930 | (346) | 91 | R 2072 | (309) | 94 |
|  | L/R entorhinal | L 556 | (98) | 112 | R 560 | (93) | 114 |
|  | L/R fusiform | L 2306 | (425) | 101 | R 2650 | (440) | 100 |
|  | L/R inferior parietal | L 4544 | (1206) | 97 | R 4825 | (1228) | 99 |
|  | L/R inferior temporal | L 3048 | (934) | 103 | R 2831 | (642) | 106 |
|  | L/R isthmus cingulate | L 1073 | (232) | 100 | R 1154 | (236) | 100 |
|  | L/R lateral occipital | L 4649 | (853) | 103 | R 4520 | (953) | 103 |
|  | L/R lateral orbitofrontal | L 2479 | (425) | 67 | R 2766 | (616) | 73 |
|  | L/R lingual | L 2528 | (566) | 100 | R 2874 | (667) | 100 |
|  | L/R medial orbitofrontal | L 1684 | (280) | 97 | R 1822 | (330) | 96 |
|  | L/R middle temporal | L 3421 | (514) | 118 | R 4045 | (965) | 111 |
|  | L/R parahippocampal | L 576 | (114) | 100 | R 579 | (84) | 100 |
|  | L/R paracentral | L 1438 | (319) | 100 | R 1444 | (396) | 100 |
|  | L/R pars opercularis | L 1315 | (130) | 98 | R 1419 | (142) | 102 |
|  | L/R pars orbitalis | L 1217 | (395) | 150 | R 1015 | (290) | 141 |
|  | L/R pars triangularis | L 1570 | (309) | 150 | R 1522 | (368) | 152 |
|  | L/R pericalcarine | L 2307 | (547) | 100 | R 2606 | (685) | 99 |
|  | L/R postcentral | L 5038 | (686) | 100 | R 4998 | (655) | 100 |
|  | L/R posterior cingulate | L 1132 | (335) | 102 | R 1301 | (501) | 101 |
|  | L/R precentral | L 4294 | (478) | 101 | R 4125 | (304) | 100 |
|  | L/R precuneus | L 3173 | (658) | 100 | R 3191 | (545) | 100 |
|  | L/R rostral anterior cingulate | L 904 | (267) | 111 | R 1121 | (317) | 138 |
|  | L/R rostral middle frontal | L 4142 | (893) | 103 | R 4066 | (961) | 103 |
|  | L/R superior frontal | L 8136 | (843) | 102 | R 8083 | (1028) | 101 |
|  | L/R superior parietal | L 4794 | (1065) | 100 | R 4590 | (1208) | 100 |
|  | L/R superior temporal | L 4401 | (805) | 118 | R 4306 | (637) | 120 |
|  | L/R supramarginal | L 3615 | (898) | 100 | R 3243 | (633) | 101 |
|  | L/R transverse temporal | L 678 | (115) | 100 | R 634 | (131) | 100 |
|  | L/R insula | L 1969 | (218) | 100 | R 1918 | (190) | 100 |

‘L/R’ indicates one label in each hemisphere. ‘S’ indicates a single label that is not separated based on hemisphere. Percentages are % of the volume of the corresponding area in the original M-CRIB atlas.

## 4 Discussion

The updated M-CRIB 2.0 atlas provides comparability with adult DKT-labelled data, and closer compatibility with subcortical segmentations derived via FreeSurfer. The parcellated images will be more readily adaptable for potential incorporation into surface-based neonatal parcellation pipelines.

Indeed, forthcoming work from our lab consists of the production of a surface-based template of the DKT-compatible cortical M-CRIB 2.0 regions, which may be utilised in combination with existing infant surface-based tools.

The individual volumetric parcellated images and T1- and T2-weighted images comprising the M-CRIB 2.0 atlas will be publicly available.

### 4.1 Conclusion

We updated the M-CRIB neonatal parcellated brain atlas to be compatible with the DKT adult cortical parcellated atlas, and to incorporate updates to subcortical regions facilitating greater compatibility with FreeSurfer’s subcortical segmentation. We achieved this via manual volumetric edits to the individual parcellated images, and via the production of a detailed, revised whole-brain parcellation protocol. The resulting M-CRIB 2.0 atlas offers greater compatibility with adult parcellated data, greater accuracy due to more reproducible landmarks, and greater optimisation for integration with surface-based infant cortical parcellation pipelines. This high-quality dataset can therefore help facilitate a broad range of investigations into brain structure and function both at the neonatal time point and developmentally across the lifespan.

